# Arthropod Abundance Associated with the Natural Resources Conservation Service: Implications for Greater Sage-Grouse Management

**DOI:** 10.1101/356451

**Authors:** H.B. Goosey, Arthropod Abundance and Grazing Livestock

## Abstract

One goal of the Natural Resource Conservation Service’s Sage-Grouse Initiative was to reverse the western US trend of declining sage-grouse populations. The sage-grouse initiative aims at preventing ‘sod-busting’ activities (conversion of native habitats into cropland) which is identified as the largest threat to stable sage-grouse populations and their habitats (USFWS 2010). Rest-rotation livestock grazing is implemented on sage-grouse ‘core areas’ with the purpose of improving rangeland health on private lands and eliminate the need of listing sage-grouse on the threatened or endangered species list. We collected arthropods in central Montana from three habitat classes associated with the Sage-grouse Initiative: 1) Grazed (actively grazed livestock pastures), 2) Deferred (Ungrazed pastures), and 3) Idle (Lands of the Lake Mason National Wildlife Refuge lower unit) and report here on 2012-2014 findings. Total arthropod catches in pitfall traps were greatest from livestock Idle pastures; however, greater numbers of those arthropods classified as sage-grouse food were caught from Deferred pastures. Differences in habitat class catches revolved primarily around the high levels of thatch found on the Lake Mason Wildlife Refuge which altered the community composition and predator:prey ratios.

## Introduction

Greater sage-grouse *(Centrocercus urophasianus;* hereafter, sage-grousej populations have been documented in decline in the western U.S. since as early as 1950s (Connelly and Bruan 1997; U.S. Fish and Wildlife Service 2010). Many factors have been highlighted as explanations of sage-grouse declines with chick and brood survival being directly linked to annual recruitment; however, the specifics of these vital rates are a poorly understood component of sage-grouse ecology (Crawford et al. 2004). Much research has been conducted on the selection criteria used by female sage-grouse when choosing a brood site (Drut et al. 1994; Sveum et al. 1998) and the results have provided a foundation for land management considerations aimed at improving sage-grouse habitat and ultimately recruitment of chicks (Connelly et al. 2004). However, the criteria that female sage-grouse use for habitat selection may not provide insight into the relationship between the site resources and chick and brood survival (Morrison 2001). The mechanisms which influence daily chick and brood survival need to be better understood and these data should be used to establish a prerequisite program which implements habitat management strategies that affect annual recruitment and, ultimately, sage-grouse populations (Gregg and Crawford 2009).

The abundance of arthropods that serve as food items of sage-grouse could be instrumental in chick survival because insects are primary foods that are critical sources of necessary nutrients (Johnson and Boyce 1990; Drut et al. 1994, Jamison et al. 2002). During the first 21-28 days post-hatch, sage-grouse chicks need to incorporate arthropods into their diets for survival and development (Johnson and Boyce 1990, Gregg et al. 2007). How food arthropod abundance, diversity, and availability vary seasonally and how they influence chick and brood survival remains little known (Gregg and Crawford 2009). However, some data suggest that the abundance of insects, particularly Lepidoptera larvae, is positively associated with survival and recruitment of sage-grouse chicks (Drut et al. 1994, Gregg and Crawford 2009). Some information is also available for Gray partridge and Sharp-tailed grouse chicks, which also feed heavily upon insects. During the first several weeks after hatching, partridge chick diet consists of 77-82% insects (Potts 1986), and growth is faster when more insects are available (Dahlgren, in Carrol 2011). This is consistent with the dependence of chick survival for other prairie-dwelling birds on insect abundance (Dreitz 2009).

Livestock grazing can alter plant communities and habitats including sites in Montana (O’Neill et al. 2003, 2010, Courtois *et al.*, 2004) which in turn either directly or indirectly influences arthropod communities (Price 2007) in a negative or positive way, depending upon the strategy of grazing. For example, Kruess and Tscharntke (2002) reported a 50% reduction in insect diversity and abundance which was attributed to the intensity and timing of the grazing event. They conclude that increased insect diversity and abundance are associated with taller vegetation which is typically located in non-grazed or strategically grazed areas. In contrast, Price (2007) suggests that moderate levels of disturbance produce the greatest arthropod abundance and diversity because disturbances open up habitats for colonists by preventing resource monopolization by competitively dominant species. From this, it is logical to suggest that a rest-rotation grazing program which leaves sufficient vegetative structure could be viewed as a moderately disturbing event with potentially beneficial influences on arthropods. However, the effect of grazing on insects depends upon both the vegetation type of the grazed habitat and the insect group considered (O’Neill et al. 2003).

We investigated, from 2012-2015, ground dwelling arthropod relative abundance and diversity in sage-brush steppe habitats found in grazed and deferred pastures associated with the Sage-grouse Initiative rest-rotation grazing system of central Montana and on the Lake Mason National Wildlife Refuge where grazing has been absent for over a decade.

The U.S. Fish and Wildlife Service (USFWS) produced a range-wide conservation report which outlines the objectives for sage grouse. The report is titled: “Greater sage grouse *(Centrocercus urophasianus)* Conservation Objectives Final Report”. The USFWS outlines conservation actions for sage-grouse management zone 1 which encompasses our research area in central Montana. The purpose of the USFWS report was to encourage conservation of sage-grouse habitats while working closely with ranch and farm owners. One of the conservation programs highlighted by the USFWS was the implementation of the Natural Resources Conservation Service (NRCS) Sage-Grouse Initiative (SGI). The SGI aims at preventing ‘sod-busting’ activities (conversion of native habitats into cropland) which is identified as the largest threat to stable sage-grouse populations and their habitats (USFWS 2010). The SGI is implemented in Montana on sage-grouse ‘core areas’ which have been determined by Montana Fish, Wildlife, and Parks (Fig. 13). The purpose of the SGI is to improve rangeland health on private lands and eliminate the need of listing sage-grouse on the threatened or endangered species list.

## Methods

### Study Area

Spring research was conducted near Lavina (46.5176 N, 108.0973), in central Montana. Smith et al. 2017 described the area as intermountain big-mountain basins big sagebrush steppe that is characterized by rolling hills with elevations of 975 m to 1,250 m. The dominant vegetation is Wyoming big sagebrush (Artemisia tridentata ssp. wyomingensis), silver sagebrush (A. cana) and a mix of perennial rhizomatous and caespitose grasses. The climate is cold and semi-arid with daily temperatures from 1981 – 2010 ranging between 2.88 C in December to 30.88 C in July with annual precipitation averaging 359 mm. During our sampling, precipitation varied from 265 mm in 2012 to 485 mm in 2014. Agricultural enterprise is the major land use and includes livestock grazing and dryland farming with approximately 10% of the area sown to crops. Smith et al. 2017 also reports that ranches that enrolled their land in NRCS SGI rotational grazing system varied in size and cattle numbers so individual grazing plans were generated to take account of these differences and individual landowner needs and that all ranch plans did adhere to the NRCS Montana Prescribed Grazing conservation practice standards. For a more detailed description of the SGI-Rotational Grazing System see Smith et al. (2017).

To direct our sampling program, we obtained the individual ranch grazing plans from each landowner which provided us with information to generate basic grazing information during our study (Table 1).

**Table 1.**
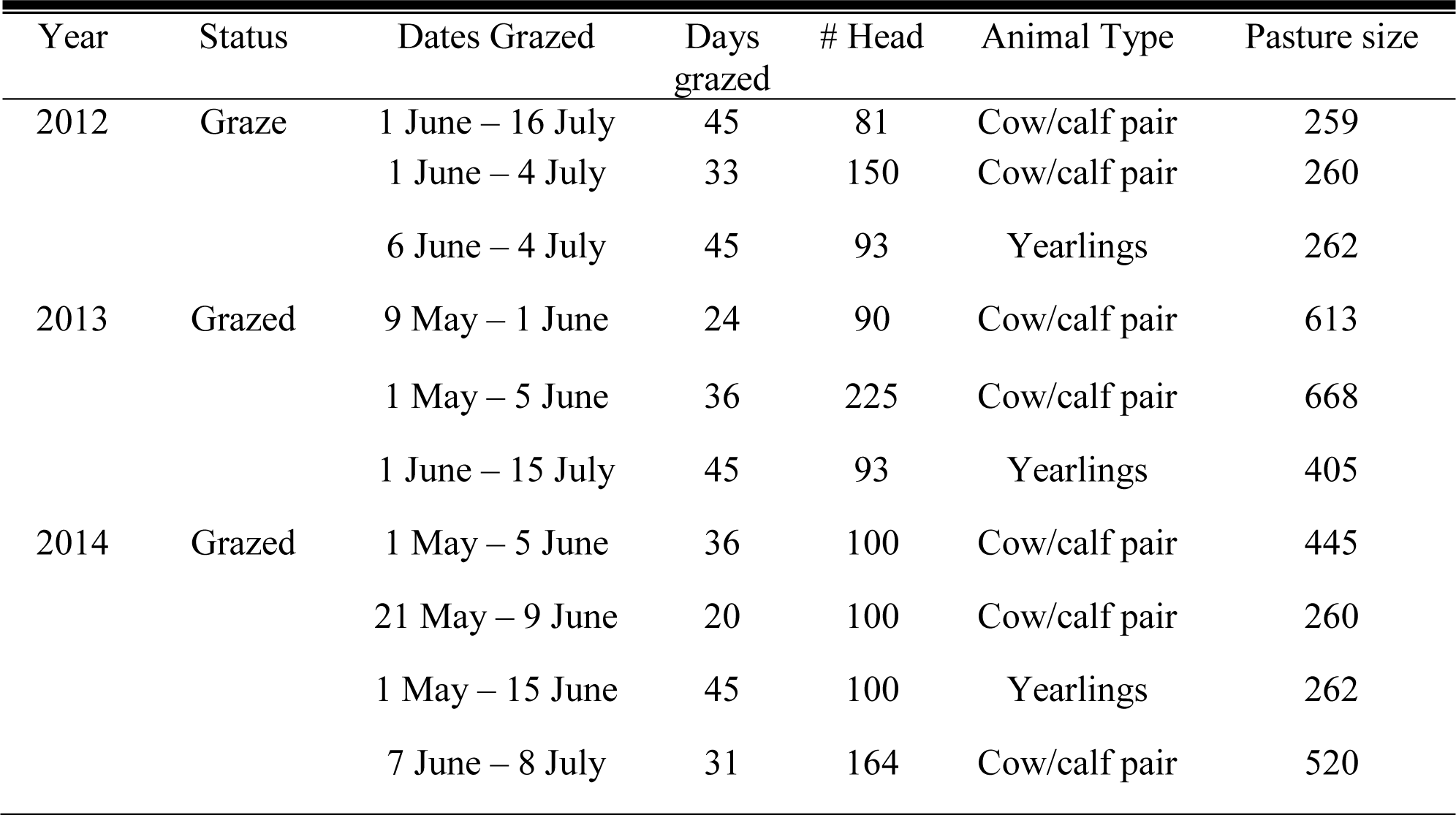
Pasture grazing plan information obtained from landowners as part of the NRCS sage-grouse initiative rest-rotation grazing program during 2012 - 2014 sampling north of Lavina, MT.

### Sampling Locations

For our research program, pastures were assigned to three treatments during our sampling periods: 1) grazed, 2) deferred, and 3) Idle. Grazed and deferred pastures were sampled during 2012 – 2014 while idle pastures were sampled during 2013 and 2014. A pasture was considered ‘Grazed’ if livestock were present during our sampling period. A pasture was considered ‘deferred’ if it was designated in the ‘Rest’ phase of the grazing program. The Lake Mason National Wildlife Refuge (LMWF) was added as an ‘Idle’ area of study and sampled in 2013 and 2014. Lands which are part of the lower unit of the LMWF have not experienced any livestock grazing for the previous 15 years (Bridget Nielson, pers. comm.).

Sampling areas in pastures were selected based on visual identification of suitable sage-grouse nesting habitat as described by Connelly et al. (2011) and exact sampling locations were randomly located within these nesting areas.

### Sampling Location Characteristics

Weekly and at each sampling location we visually estimated the percentage of bare ground by placing a 0.5 m^2^ metal ring 2 m apart along a random compass bearing originating from the center of our pitfall trap transect. Along the same random compass bearing, we measured the height of live grass and live sagebrush at 10 random locations.

### Arthropod Sampling

The activity density of ground-dwelling arthropods was assessed in each experimental unit by deploying 10 pitfall traps, measuring 9cm wide, 1m apart on a linear transect which was determined by a random compass. To construct the pitfall traps, we dug ~20-30cm-deep X 10-cm-wide holes with a post-hole auger and placed two stacked 0.5-liter plastic cups (Solo Cup Company, Lake Forest, IL) in each of those holes. We backfilled the pitfall trap holes until the mouth of the top cup was flush with the soil surface and filled the top cup of the pitfall trap approximately one-third full of propylene glycol-based antifreeze (Arctic Ban, Camco Manufacturing Inc., Greensboro, NC). Each pitfall trap was covered with a rain cover constructed from a 25-cm-diameter clear plastic plate held to the ground with three equally spaced 10-cm bolts. All rain covers had at least 2 cm between the soil surface and the rim of the clear plastic plates to avoid interfering with ground dwelling arthropod activity. Traps were deployed in sage grouse nesting/brood rearing habitat during the period of late nesting to late brooding to capture arthropods available for chick consumption during this time. While installed, each week we collected all arthropods caught in the pitfall traps by placing them in an 11.5 × 23cm plastic bag (Whirl-Pak, Nasco Inc., Fort Atkinson, WI). Following collection, all pitfall traps were replenished with antifreeze for sampling the subsequent week. Traps were deployed and actively collecting specimens 24 hrs. per day for the entire sampling period. We sorted all samples for Coleoptera, Hymenoptera (Formicidae), Orthoptera, Lepidoptera (larvae), and Aracnida, transferred them to 70% by volume ethanol and identified them to family in the laboratory following (Triplehorn and Johnson 2005).

One limitation of pitfall trapping is that the probability of capturing a specimen depends on both how many beetles are in each area and how much those specimens are moving (Luff 2002). Thus, pitfall trapping confounds activity and density. Entomologists, therefore, refer to values obtained from pitfall trapping as “activity-density,” and treat those values as metrics of relative abundance (Kromp 1989). Additionally, because in our study beetles were free to disperse between fields, our results reflect habitat selection rather than changes in carabid populations per se (Lee et al. 2001). While these limitations have been well documented in the literature, pitfall trapping remains one of the most efficient methods for sampling ground swelling arthropds (Greenslade 1964, Kromp 1989, Spence and Niemela” 1994, Lee et al. 2001, Luff 2002).

### Statistical Analysis

Regression function linearity was evaluated by plotting the independent (X) data versus the residuals. Treatment homogeneity of variances were estimated by plotting predicted values versus residuals and by analyzing the absolute values of the residuals and location as a class variable (Proc Mixed, SAS 2008). Error term independence was evaluated by plotting the residuals versus time.

Catch data were organized by functional group analyses of 1) Predators (i.e., Carabidae, Lycosidae, Gnaphosidae, Philodromidae, Salticidae, Thomisidae, and Hahniidae), Detritivores (i.e., Tenebrionidae, Scarabaeidae, Histeridae, Dermestidae, Silphidae, and Nitidulidae), and Food Arthropods (i.e., Carabidae, Tenebrionidae, Scarabaeidae, Chrysomelidae, Curculionidae, Coccinellidae, Arctiidae, Saturniidae, Pieridae, Formicidae, Gryllidae, Acrididae, Tettigoniidae). Total arthropod catches were also analyzed.

Analyses were carried out on data that was summed across all eight sampling dates and summed again over three years of study in SGI pastures and two years of study at the LMWR for a total multi-year catch associated with sampling locations. Multiple comparison data were analyzed using the Proc Mixed procedures SAS^®^ version 9.2 for Multi-Location Data with pasture as the experimental unit. Sampling locations were considered random and nested within year with the Kenward-Roger option used to determine degrees of freedom, which is the recommended and most conservative option for unbalanced data sets (Littell et al. 2002). Locations effects are considered random if the locations plausibly represent the population (Littell et al. 2002). The sample size equation 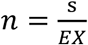 (Southwood and Henderson 2000) was used to calculate the number of necessary replicates.

A second order polynomial equation *y* = *ax*^2^ + *bx* + *c* was generated using SigmaPlot^®^ v. 11.2 (SigmaPlot^®^ 2009) where *x* is bare ground, which was visually estimated at each sampling location and where, *a, b, and c* are regression coefficients and where *y* is food arthropod catches.

To further analyze the overall similarities among our sampling locations within Deferred, Grazed, and Idle sampling areas, we used PAST v. 3.19 (Hammer et al. 2001) to carry out a detrended correspondence analysis (DCA) on location by taxa abundance matrices for each location and year. All arthropod specimens were identified from each sample and included in the DCA analyses.

### Results

Specimens were collected and identified during 3 years of study. Regression linearity, independence, and error term normality were met. Variance homogeneity was met across location *(F* = 2.46; df = 3, 22; *P* = 0.0895) and treatment (F = 1.49; df = 2,23; *P* = 0.2471).

Treatment x year interactions did not differ for weekly total arthropod catches (*F* = 1.10; df = 3, 25; *P* = 0.3759), total predators (*F* = 0.38; df = 3, 25; *P* = 0.7676), total detritivores (*F* = 0.04; df = 3, 25; *P* = 0.9887), and total Lepidoptera larvae (*F* = 3.26; df = 2, 25; *P* = 0.06), and total food arthropods (*F* =1.95; df = 3, 25; *P* = 0.1580), therefore data were combined across year for analyses.

Treatment total arthropod catches with more (*F* = 10.59, df = 2, 14; *P* < 0.01) captures from LMWR Idle than SGI Grazed or Deferred (Fig. 1a). Greater numbers of predators (*F* = 26.49; df = 2, 14; *P* <0.01) and detritivores (*F* = 10.77; df = 2, 14; *P* < 0.01) were captured from LMWF Idle than in SGI Grazed or Deferred pastures (Fig. 1b, 1c). Fewer Lepidoptera larvae (*F* = 8.54; df = 2, 14; *P* <0.01) were captured from LMWR Idle than from SGI Grazed and Deferred pastures (Fig. 1d). Sage-grouse Food Arthropods catches differed with more captures from Deferred than from Idle (t = 2.44; df = 2, 13.9; *P =* 0.0504) and no differences in captures among Deferred and Grazed (t = 0.49; df = 2, 13.4; *P* = 0.6306) or among Idle and Grazed (t = 1.73; df = 2,13.9; *P =* 0.1048).

## Discussion

The lack of livestock grazing at the Lake Mason Wildlife Refuge has resulted in an accumulation of plant litter which can have a cascading effect on the microclimate, predator ambush sites and ultimately insect diversity. O’Neill et al. 2003 excluding livestock in south west Montana for more than six years period from experimental plots of crested wheat grass. The long term exclosures unintentionally copied some of the effects of CRP by increasing crested wheat grass predominance. Without livestock grazing, plant litter accumulated and densities of grasshoppers, and black grass bugs increased and a microclimate modification affected thermoregulatory abilities of grasshoppers (O’Neill et al. 2003, 2008, 2010, O’Neill and Rolston 2007).

Further discussions are forthcoming.

## References Cited

Campbell, G. L., A. A. Marfini, R. S. Lanciotti, and D. J. Gubler. 2002. West Nile virus. Lancet: Infectious Disease. 2: 519–529.

Connelly, J. W., and E. C. Braun. 1997. Long-term changes in sage grouse *Centrocercus urophasianus* populations in western North America. Wildlife Biology. 3: 229–234.

Connelly, J. W. S. T. Knick, M. A. Schroeder, and S. J. Stiver. 2004. Conservation assessment of greater sage-grouse and sagebrush habitats. Western Association of Fish and Wildlife Agencies unpublished report. Cheyenne, Wyoming. USA.

Connelly, J. W., E. T. Rinkes, and C. E. Braun. 2011. Characteristics of greater sage-grouse habitats: a landscape species at micro-andmacroscales. Pp. 69–83 In S.T. Knick and J. W. Connelly (eds). Greater Sage-Grouse: ecology and conservation of a landscape species and its habitats. Studies in Avian Biology vol. 38, University of California Press, Berkeley, CA.

Connelly, J. W., M. A. Schroeder, A. R. Sands, and C. E. Braun. 2000. Guidelines to manage Sage Grouse populations and their habitats. Wildlife Society Bulletin. 28: 967–985.

Courtois, D. R., B. L. Perryman, and H. S. Hussein. 2004. Vegetation change after 65 years of grazing and grazing exclusion. J. Range Management. 57: 574–582.

Crawford, J. A., R. A. Olson, N. E. West, J.C C. Mosley, M. A. Schroeder, T. D. Whitson, R. F. Miller, M. A. Gregg, and C. S. Boyd. 2004. Ecology and management of sage-grouse and sage-grouse habitat. J. Range Management. 57: 2–19.

Dahlgren, J. 1990. The significance of arthropods in the gray partridge diet, pp. 202–213. In Perdix V: Gray Partridge and Ring-necked Pheasant Workshop. Kans. Dep. Wildl. Parks, Emporia, KS.

Doherty, K. E. 2008. Sage-grouse and energy development: integrating science with conservation planning to reduce impacts. Ph.D. dissertation, University of Montana, Missoula, MT.

Dreitz, V. J. 2009. Parental behaviour of a precocial species: implications for juvenile survival. Journal of Applied Ecology. 46: 870–878.

Drut, M. S., W. H. Pyle, and J. A. Crawford. 1994. Diets and food selection of sage grouse chicks in Oregon. J. Range Management. 47: 90–93.

Gaines, H. R. and C. Gratton. 2010. Seed predation increases with ground beetle diversity in a Wisconsin (USA) potato agroecosystems. Agriculture, Ecosystems, and Environment. 137: 329–336.

Gregg, M. A., and J. A. Crawford. 2009. Survival of greater sage-grouse chicks and broods in the northern great basin. J. Wildlife Management. 73: 904–913.

Gregg, M. A., M. R. Dunbar, and J. A. Crawford. 2007. Use of implanted radio transmitters to estimate survival of greater sage-grouse chicks. J. Wildlife Management. 71: 646–651.

Jamison, B. E., R. J. Robel, J. S. Pontius, and R. D. Applegate. 2002. Invertebrate biomass: associations with lesser prairie-chicken habitat use and sand sagebrush density in southern Kansas. Wildlife Society Bulletin. 30: 517–226.

Johnson, G. D., and M. S. Boyce. 1990. Feeding trials with insects in the diet of sage grouse chicks. J. Wildlife management. 54: 89–91.

Kruess, A., and T. Tscharntke. 2002. Contrasting responses of plant and insect diversity to variation in grazing intensity. Biological Conservation. 106: 293–302.

Lukacs, P. M., A. B. Franklin, and D. R. Anderson. 2004. Passive approaches to detection in distance sampling, pp. 260–280. *In* S. T. Buckland, D. R. Anderson, K. P. Burnham, J. L. Laake, D. L. Borchers, and L. Thomas (eds.) Advanced distance sampling: estimating abundance of biological populations. Oxford University Press, New York, NY.

Morrison, M. L. 2001. A proposed research emphasis to overcome the limits of wildlife-habitat relationship studies. J. Wildlife Management. 65: 613–623.

NatureServe. 2016. NatureServe Explorer: an online encyclopedia of life [web application]. NatureServe, Arlington, Virginia, USA. http://explorer.natureserve.org

Naugle, D. E., C. L. Aldridge, B. L. Walker, T. E. Cornish, B. J. Moynahan, M. J. Holloran, K. Brown, G. D. Johnson, E. T. Schmidtmann, R. T. Mayer, C. Y. Kato, M. R. Matchett, T. J. Christiansen, W.E. Cook, T. Creekmore, R. D. Falise, E. T. Rinkes, and M. S. Boyce. 2004. West Nile virus: pending crisis for Greater Sage-Grouse. Ecology Letters. 7: 704–713.

O’Neill K.M., and M.G. Rolston. 2007. Short-term dynamics of behavioral thermoregulation by adults of the grasshopper *Melanoplus sanguinipes*. 14pp. J. Insect Sci. 7:27, available online: insectscience.org/7.27.

O’Neill, K.M., B.E. Olson, M.G. Rolston, R. Wallander, D.P. Larson, and C.E. Seibert. 2003. Effects of livestock grazing on rangeland grasshopper (Orthoptera: Acrididae) abundance. Agric. Ecosyst. Environ. 97: 51–64.

O’Neill, K.M., B.E. Olson, R. Wallander, M.G. Rolston, D.P. Larson, and C.E. Seibert. 2010. Effects of livestock grazing on grasshopper abundance on a native rangeland in Montana. Environ. Entomol. 39: 775–786.

O’Neill, K.M., S. Blodgett, B.E. Olson, and R.S. Miller. 2008. Black grass bug (Hemiptera: Miridae; *Labops hesperius* Uhler, *Irbisia* spp.) abundance correlates with grazing intensity in crested wheatgrass (*Agropyron cristatum* (L.) Gaertn.) pastures in Montana. J. Econ. Entomol. 101: 309–313.

Potts, G. R. 1986. The partridge: pesticides, predation, and conservation. Collins: London xiii + 274 pp.

Price, P. W. 1997. Diversity and Stability, pp. 659–685. *In* Price, P. W. (eds.) Insect ecology, 3^rd^ ed. John Wiley & Sons, Inc., New York, NY.

SAS Institute Inc. 2008. SAS/STAT^®^ 9.2 User’s Guide. Cary, NC: SAS Institute Inc

SigmaPlot. 2009. SigmaPlot^®^ 11.2 User’s Guide. San Jose, CA: Systat Software, Inc.

Smith. J.T., J.D. Tack, L.I. Berkeley, M. Szczypinski, and D. E. Naugle. 2017. Effects of Rotational Grazing Management on Nesting Greater Sage-Grouse. J. Wildlife Management. DOI: 10.1002/jwmg.21344

Southwood, T. R. E., and P. A. Henderson. 2000. The sampling programme and the measurement and description of dispersion, pp. 7–72. *In* T. R. E. Southwood and P. A. Henderson (eds.) Ecological methods, 3^rd^ ed. Blackwell Science Ltd, Oxford, UK.

Sveum, C. M., J. A. Crawford, and W. D. Edge. 1998. Use and selection of brood-rearing habitat by sage grouse in south-central Washington. Great Basin Naturalist. 58: 344–351.

Triplehorn, C. A. and N. F. Johnson. 2005. Study of insects. 7^th^ edition.

U.S. Fish and Wildlife Service. 2010. Endangered and threatened wildlife and plants; 12-month finding for petitions to list the greater sage-grouse (*Centrocercus urophasianus*) as threatened or endangered. Federal Register 75: 13910–14014.

